# The Speciation Continuum in Bloom: Incomplete Lineage Sorting, Gene Flow, and Reticulate Evolution in Rapidly Diverging Plant Lineages

**DOI:** 10.64898/2026.03.30.715067

**Authors:** Luana S. Soares, Nelson J. R. Fagundes, Aureliano Bombarely, Loreta B. Freitas

## Abstract

The remarkable diversity of life on Earth results from evolutionary processes functioning across different spatial and temporal scales. Species diversification occurs through various mechanisms and at widely varying rates, but identifying the conditions that trigger bursts of diversification over short timescales remains a central challenge in evolutionary biology. This difficulty is more pronounced when incomplete lineage sorting (ILS), hybridization, and ongoing gene flow obscure evolutionary relationships and complicate species delimitation. In this study, we investigated the evolutionary history and species boundaries within a group of recently diverged *Petunia* lineages shaped by pervasive gene flow. We integrated phylogenomic, population genetic, and species delimitation approaches to reconstruct lineage relationships and assess whether these lineages represent distinct species or stages along a speciation continuum. By applying methods that account for both ILS and gene flow, we revealed that most lineages are not fully independent evolutionary units but rather occupy intermediate positions along this continuum. Gene flow played a crucial role during diversification, blurring species boundaries and generating reticulate evolutionary patterns. Our findings demonstrate that traditional phylogenetic trees may oversimplify relationships in such systems, while phylogenetic networks offer a more accurate representation of evolutionary history. Comprehensive and integrative analyses, such as those employed here, are essential for capturing these complex dynamics. Ultimately, only four lineages could be confidently recognized as distinct species, whereas the remaining represent cases of ongoing divergence. These results emphasize the need to refine species delimitation frameworks for systems characterized by recent divergence and extensive reticulation.

## Introduction

The extraordinary diversity of life on Earth stems from countless evolutionary processes operating across spatial and temporal scales. Speciation, the formation of new, independently evolving lineages (de Queiroz 2007), is fundamental to generating and maintaining biodiversity. Speciation occurs when diverging populations accumulate ecological, functional, and/or reproductive differences that ultimately prevent them from merging into a single evolutionary unit (Feder et al. 2012; Nosil and Feder 2012). However, the timescale of speciation can vary widely among taxa (Stebbins 1974; Stenseth and Smith 1984), and the factors underlying rate variation, particularly those that enable bursts of diversification over short timescales, remain a key and unresolved issue in evolutionary biology (Losos and Schluter 2000; Hughes and Eastwood 2006).

Despite significant advances in molecular and genomic approaches, species delimitation still faces conceptual and operational challenges (de Queiroz 2007; Carstens et al. 2013; Rannala 2015; Sukumaran and Knowles 2017; Smith and Carstens 2020). Much of this difficulty stems from a lack of consensus on what defines a species (Sukumaran and Knowles 2017). Criteria for species delimitation often overlap with species concepts, and disagreements about conceptual frameworks prevent the development of universally applicable, objective methods for species identification (Hey 2001; Carstens et al. 2013; Rannala 2015). Criteria such as distinguishable morphology or intrinsic reproductive barriers do not arise simultaneously, thereby challenging the recognition of independent evolutionary lineages in the so-called “grey zone” of speciation (de Queiroz 2007; Roux et al. 2016).

Delimiting species is particularly challenging in cases of recent and rapid divergence, where factors such as incomplete lineage sorting (ILS), hybridization, and ongoing gene flow obscure evolutionary relationships (Jackson et al. 2017; Meier et al. 2017; Barley et al. 2024). In these systems, diverging lineages often remain interconnected through gene flow for extended periods.

Depending on the selective pressures on external morphology, this can result in either phenotypically distinct yet genetically entangled populations (Dufresnes et al. 2020, 2023; Chambers et al. 2023; Barley et al. 2024) or, conversely, in morphologically undistinguished but genetically differentiated species (Pfenninger and Schwenk 2007; Struck et al. 2018; Soares et al. 2024). These complexities are especially pronounced in plants, where hybridization is common. For example, at least 25% of the flora in the United Kingdom actively hybridizes with relatives (Mallet 2005; Stace et al. 2015), and 40-70% of living plant species are polyploids, with ca. 15% of angiosperm species being allopolyploids (Doyle and Sherman-Broyles 2017). In phylogenomic studies, recent radiations introduce additional challenges: rapid divergence leads to short internal branches, amplifies ILS, and heightens the likelihood of introgression, all of which complicate species tree estimation and the delineation of species boundaries (Abbott et al. 2013; Eaton and Ree 2013; Malinsky et al. 2018; Steenwyk et al. 2023).

*Petunia* Juss. (Solanaceae) serves as an ideal system for investigating the genomic signatures of speciation in the presence of gene flow within a context of recent and rapid divergence. Native to the lowlands and highlands of southern South America, this group of annual herbaceous plants comprises 20 recognized species (Soares et al. 2025) that have recently diverged during the climatic fluctuations of the Pleistocene (Fregonezi et al. 2013; Freitas 2022; Soares et al. 2023; 2024; Pezzi et al. 2024). These climatic oscillations shaped species distributions and likely fostered cycles of isolation and secondary contact, making *Petunia* an excellent case study for examining the interplay between divergence, introgression, and incomplete lineage sorting.

Although most research has focused on lowland species (Soares et al. 2025), highland species remain comparatively underexplored. These taxa, which are restricted to the highland regions of southern Brazil and neighboring areas, exhibit unsolved taxonomic boundaries and conflicting phylogenetic relationships (Lorenz-Lemke et al. 2010; Souza et al. 2022; 2024; Soares et al. 2023; 2024; Soares and Freitas 2024). Therefore, understanding the evolutionary history of these highland lineages is crucial for reconstructing the diversification dynamics of the entire genus and clarifying the process of speciation.

This study aimed to examine the species delimitation boundaries within a recently divergent group of lineages, unveiling their evolutionary history shaped by pervasive gene flow. We combined phylogenomic, population genetic, and species delimitation approaches to reconstruct lineage relationships, estimate the extent of introgression, and evaluate the continuity of the speciation process. By integrating methods that account for both incomplete lineage sorting and gene flow, we sought to determine whether these lineages represent distinct species or points along a speciation continuum. Ultimately, our goal was to clarify the limitations of traditional bifurcating phylogenies in reticulate systems and to highlight the importance of network-based frameworks for representing evolutionary history in recent, dynamic radiations.

## Materials & Methods

### Sampling, DNA Extraction, and Sequencing

We sampled young and healthy leaves from 132 individuals across 11 *Petunia* species with short corolla tubes (Table S1). Based on previous results (Soares et al. 2024; 2025), the species *P. guarapuavensis* (Pgua), *P. scheideana* (Psch), and *P. interior* (Pteri) were subdivided into their respective intraspecific lineages (Fig. 1, Fig. 2a). In certain analyses, we used *P. axillaris* as an outgroup because this species belongs to a different *Petunia* clade (Reck-Kortmann et al. 2014; Soares et al. 2025). The sample size per collection site adhered to the guidelines for non-model species in population genomic analyses (Nazareno et al. 2017) and was comparable to studies involving *Petunia* species (Caballero-Villalobos et al. 2021; Giudicelli et al. 2022; Guzmán et al. 2022; Pezzi et al. 2022; Soares and Freitas 2024). Vouchers were collected from each population and deposited at the herbarium of the Universidade Federal de Minas Gerais (BHCB-UFMG) in Belo Horizonte, Brazil.

**Figure 1.**
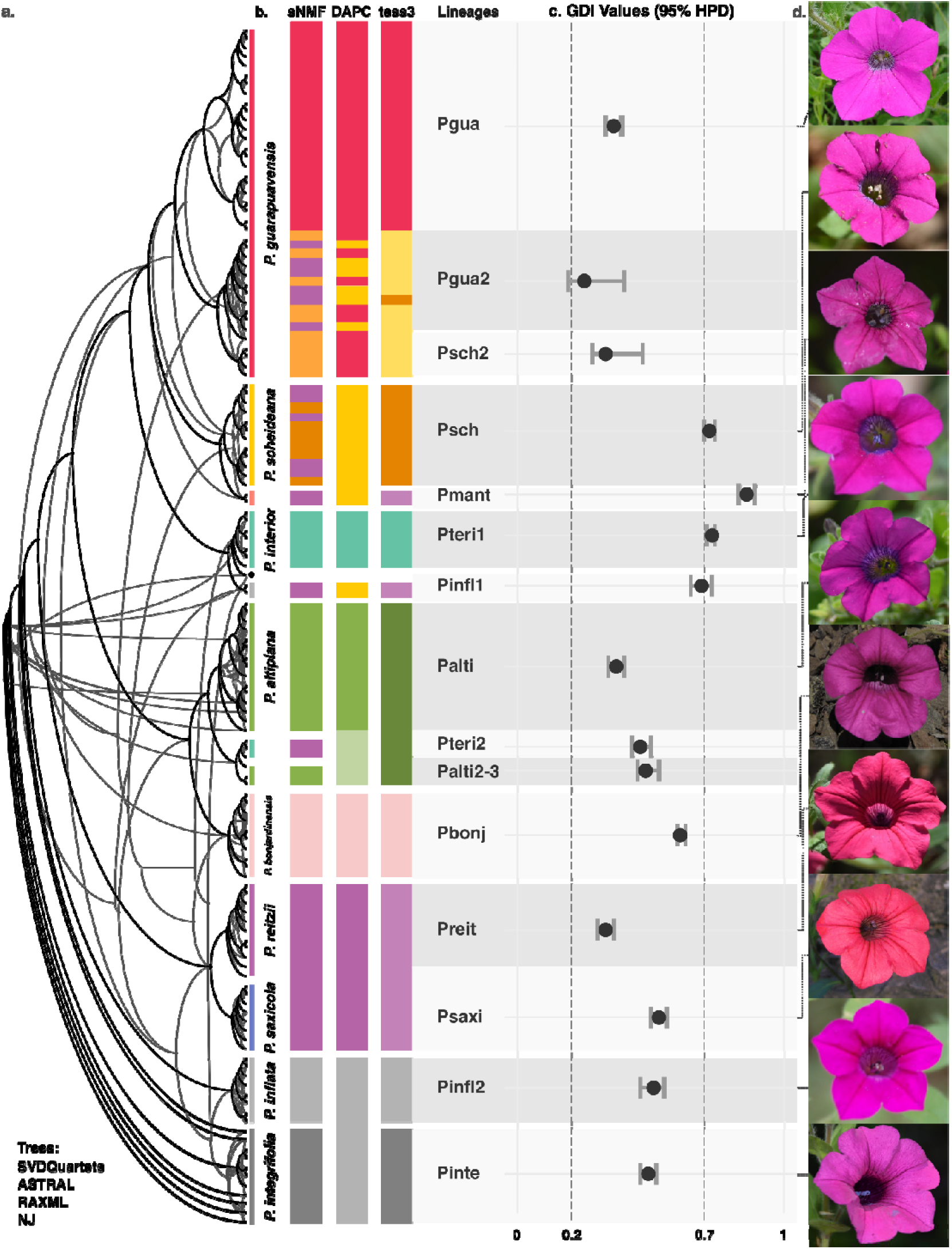
Evolutionary relationships, genetic structure, and morphological variation among highland *Petunia* species. (a) Phylogenetic relationships inferred using four different approaches: SVDQuartets, ASTRAL, RAxML, and Neighbor-Joining (NJ), visualized as a DensiTree. Species and lineages are color-coded consistently across analyses. (b) Population structure results showing the most likely population assignment for each individual based on the highest ancestry proportion (Q-value) from three different clustering methods: sNMF, DAPC, and TESS3. (c) Genealogical divergence index (GDI) values were estimated for each lineage, indicating levels of genetic differentiation and the support for species delimitation. (d) Representative floral morphologies for some of the species and lineages included in this study.

**Figure 2.**
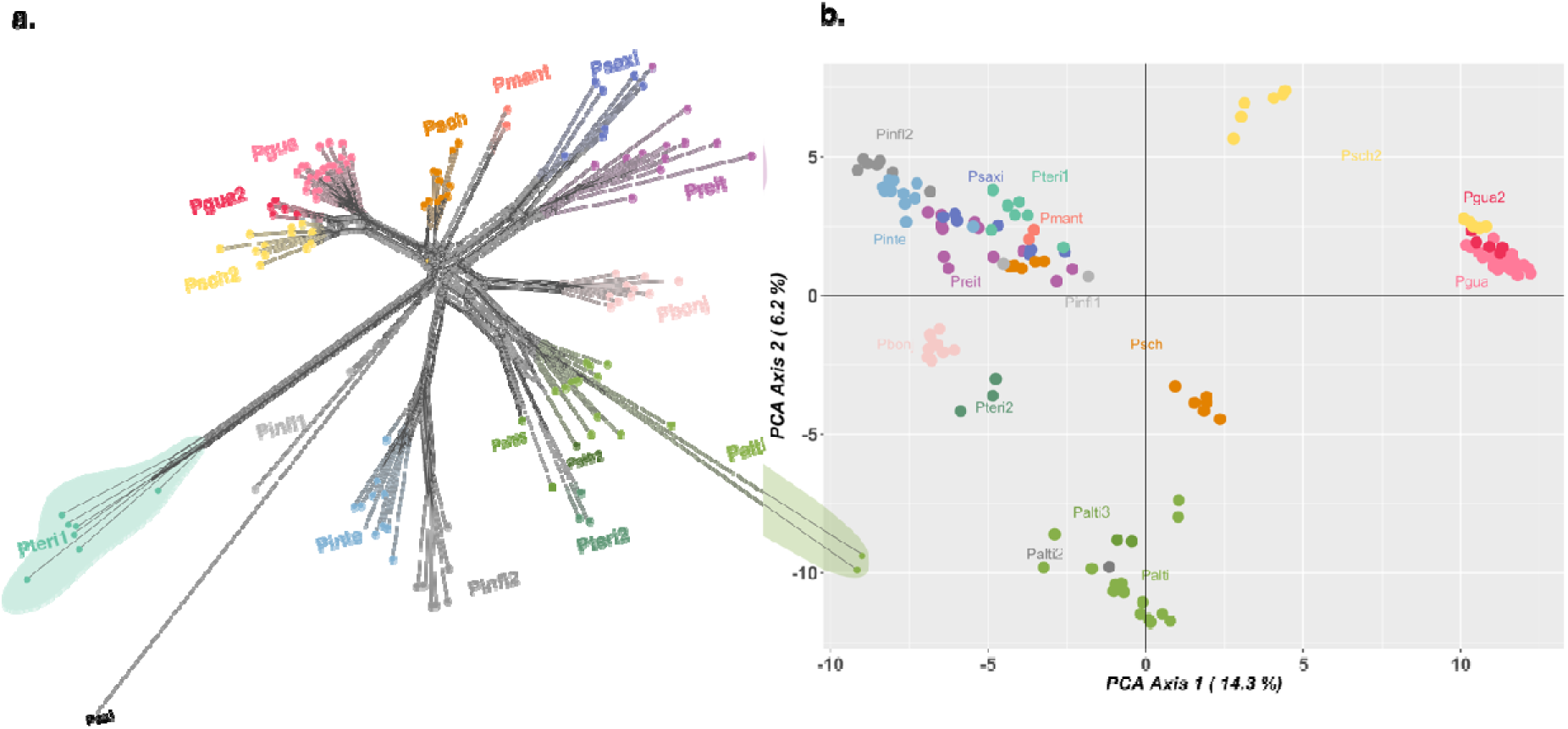
(a) Principal Component Analysis (PCA) based on SNP data, showing the genetic structure among lineages. Colors correspond to predefined lineage groups used throughout the study. (b) Phylogenetic network inferred with SPLITSTREE, illustrating the relationships among individuals and lineages. Lineages are color-coded consistently across all panels.

We powdered the silica-dried leaves with liquid nitrogen to extract genomic DNA using a cetyltrimethylammonium bromide (CTAB; Sigma-Aldrich Chem. Co., St. Louis, USA) protocol (Roy et al. 1992). We measured DNA concentration using a Qubit Fluorometer (Thermo Fisher Scientific Co., Waltham, USA) and assessed DNA quality using a NanoDrop DN-1000 Spectrophotometer (Thermo Fisher). DNA samples with a 260/280 ratio > 1.80 and a 260/230 ratio > 1.80 were regarded as high quality and used to construct the genomic library.

We processed the DNA library with DARTSEQ^TM^ (Kilian et al. 2012; Cruz et al. 2013) to reduce complexity using a method that combines the *Pst*I and *Mse*I enzymes (NEB - New England BioLabs Inc., Ipswich, USA) and replaces a single adaptor with two (Kilian et al. 2012).

Sequencing involved combining equimolar amounts of amplification products from each 96-well microtiter plate sample and using them in a c-Bot bridge PCR system (Illumina Inc., San Diego, USA), followed by sequencing on the Illumina HiSeq 2500 platform (Illumina).

### Filtering and Variant Discovery

We inspected the raw data using FASTQC v.0.11.7 (Andrews, 2010) and MULTIQC (Ewels et al. 2016). Additionally, we removed barcodes and adapters, trimmed low-quality regions (< Q30), and discarded reads shorter than 50 bases using FASTQ-MCF v.1.04.807 (Aronesty 2013). The reads were mapped against the *P. inflata* v.1.0.1 reference genome (Bombarely et al. 2016) using BWA v.0.7.10-r789 (Li and Durbin 2010) with default settings. Unmapped reads were removed, and individual SAM files were converted to BAM format using SAMTOOLS v.1.3.1 (Danecek et al. 2021). All BAM files were merged into a single file using the *bamaddrg* utility (available at https://github.com/ekg/bamaddrg), then sorted and indexed using SAMTOOLS.

We used FREEBAYES v.1.3.6 (Garrison and Marth 2012) to call variants by applying the following parameters: mapping quality > 30, base quality > 30, and read depth > 10. We ran VCFTOOLS v.0.1.12 (Danecek et al. 2011) to filter and retain only biallelic single-nucleotide polymorphisms (SNPs) with up to 10% missing data and a minimum allele frequency (--maf) of 0.01. To remove loci in linkage disequilibrium (LD), we set the minimum site distance to 100 bp (--thin) and kept only one SNP per read. Outlier loci were detected using PCADAPT (Luu et al. 2017) and removed for demographic analyses. When necessary, the raw VCF file was subdivided using BCFTOOLS v.1.13 (Danecek et al. 2021) and filtered again with the new set of individuals. The VCF file was converted to different formats using the PGDSPIDER v.2.1.1.2 (Lischer and Excoffier 2012) and DARTR v.2.05 (Mijangos et al. 2022) R packages to perform further analyses.

### Evolutionary Relationships

To investigate evolutionary relationships among species, we employed multiple phylogenetic approaches. First, we constructed a relationship network using the NeighborNet method in SPLITSTREE v.4.16 (Huson and Bryant 2006). We then inferred several phylogenetic relationships using Neighbor Joining (NJ), maximum likelihood (ML), SVDQUARTETS, SNAPP, ASTRAL with RAxML trees, and IQ-TREE, with *P. axillaris* as the outgroup. The NJ analysis was conducted with the *gl.tree.nj* function of the DARTR R package. For the ML analysis, we generated a species tree using RAxML-NG v.1.0.0 (Kozlov et al. 2019), applying the GTR+ASC_LEWIS model with 1,600 bootstrap replicates. The VCF file containing the SNP matrix was converted into PHYLIP format using PGDSPIDER v.2.1.1.2 (Lischer and Excoffier 2012), and these ML trees were used to perform MSC analyses in ASTRAL v.5.7.8 (Zhang et al. 2018).

We also employed the coalescent-based SVDQUARTETS method (Chifman and Kubatko 2014), implemented in PAUP* v.4a (Swofford 2015). We evaluated all possible quartets through 100 bootstrap replicates to generate a 50% majority-rule consensus tree. In parallel, we constructed a maximum likelihood phylogeny using IQ-TREE v.2.3.6 (Minh et al. 2020), estimating branch support with 1000 ultrafast bootstrap replicates (UFB; Hoang et al. 2018).

To estimate lineage divergence times, we ran SNAPP (Bryant et al. 2012), a method based on the multispecies coalescent model for SNP data. For calibration, we constrained the crown divergence between *Petunia* and *Calibrachoa* Cerv., a genus evolutionarily close to *Petunia* (Reck-Kortmann et al. 2014), using a dated phylogeny (Särkinen et al. 2013). The estimated age of this node was set to 2.85 Ma (95% HPD: 5.5–11.67 Ma) and was modeled as a *log*-normal distribution centered at 2.85 Ma with a standard deviation of 0.16, as calculated using BEAUTI from BEAST v.1.7 package (Drummond et al. 2012). We prepared the input data using the *snapp_prep.rb* script (Stange et al. 2018), limiting the dataset to 1000 randomly selected SNPs and setting the MCMC chain length to 100,000 iterations. Two independent analyses were performed, and *log* files and trees were combined using LOGCOMBINER v.2.7.7, part of BEAST v.2.7.7 (Bouckaert et al. 2014). We assessed convergence (ESS > 200) using TRACER v.1.6 (Rambaut et al. 2018) and visualized tree topologies and node heights with DENSITREE (Bouckaert 2010) and FIGTREE v.1.4.4 (available at https://github.com/rambaut/figtree/).

Incongruences within the trees were quantified using the generalized Robinson-Foulds metric, following Smith (2020) and utilizing the TREEDIST R package (available at https://github.com/ms609/TreeDist). Full concordance was indicated by a value of 0, whereas full discordance was indicated by a value of 1.

### Genetic Diversity and Lineage Structuration

We used the filtered dataset for all analyses, excluding monomorphic sites (variants appearing in only one individual compared to the reference genome). We conducted a principal components analysis (PCA) using the *gl.pcoa* function in DARTR.

The lineage structure was determined using a Discriminant Analysis of Principal Components (DAPC; Jombart et al. 2010) in the ADEGENET v.2.1.3 R package (Jombart 2008), using the *find.clusters* and *optim.a.score* functions to identify the optimal number of *K* genetic groups. Population structure was further assessed using the SNMF software (Frichot et al. 2014), with 10 runs for each *K* value ranging from 2 to 16. The best-fit *K* was determined based on the lowest cross-entropy criterion, which was calculated using the *sNMF* function in the LEA R package.

To evaluate genetic structure in a spatial context, we used TESS3 (Caye et al. 2016) and incorporated geographic coordinates into the analysis. Ancestry maps were created using the GGPLOT2, MAPS, and RASTER v.4.2.0 R packages. We computed the most likely population assignment for each individual based on the highest ancestry proportion (*Q*-value) from three different clustering methods.

Finally, we calculated the pairwise fixation index (*F*_ST_) in ARLEQUIN v.3.5.2.2 (Excoffier and Lischer 2010) to quantify genetic differentiation between lineages, and we employed STACKS v.2.41 (Rivera-Colón and Catchen 2022) to obtain the nucleotide diversity (π, calculated per site), inbreeding coefficients (*F*_IS_), and private alleles.

### Introgression/Gene Flow Analyses

We employed complementary analyses that account for incomplete lineage sorting to investigate introgression. We assessed signatures of introgression using the asymmetric four-taxon (ABBA-BABA) and the symmetric five-taxon (DFOIL) tests based on previously obtained phylogenomic results. The ABBA-BABA (Green et al. 2010; Durand et al. 2011) analysis was conducted with DSUITE (Malinsky et al. 2021), integrating *f*-branch statistics (Malinsky et al. 2018) to visualize shared introgression across the tree. *Z*-scores for *f*-branch values were calculated using the -Z argument, and results were summarized with the *dtools.py* script. To further examine gene flow, we performed DFOIL (Pease and Hahn 2015) using EXDFOIL (Lambert et al. 2019), which extends *D*-statistics into a symmetric five-taxon framework, allowing us to infer the direction and timing of introgression with high accuracy. Additionally, we used *D*-statistics in HYDE (Blischak et al. 2018) to test for putative hybrid origins of the lineages, using a *P*-value cutoff of 0.05.

Given the evidence of admixture between the taxa from the initial analyses (see Results), we estimated their evolutionary history while accounting for introgression. We searched for networks using SNAQ (Solís-Lemus and Ané 2016), implemented in the PHYLONETWORKS v.0.12.016 package. SNAQ is a pseudolikelihood method that estimates a phylogenetic network while considering ILS and gene flow. We used the script *SNPs2CF* (available at https://github.com/melisaolave/SNPs2CF) to estimate the quartet-level concordance factor (*quartetCF*) from the VCF file with resampling for bootstrapping. The *quartetCFs* were used to infer maximum pseudolikelihood (MPL) multispecies coalescent networks (MSC-N) with the *snaq!* algorithm of PHYLONETWORKS. First, one MSC-N with zero reticulations (essentially a tree) was inferred with the *quartetCFs*, using the unrooted tree from *SVDquartets* as a starting point. For the subsequent MSC-N estimations, with 1 to 8 reticulations (*hmax*), we used the first MSC-N (*hmax*L=L0) as input. The best number of hybridizations was determined by plotting the *log*-likelihoods from the runs against the *hmax,* following the recommendation in the online manual.

Additionally, we used ADMIXTOOLS2 in R (available at https://uqrmaie1.github.io/admixtools) to estimate admixture graphs. The best-fit graphs were estimated using 100 independent runs of the *find_graphs command* (stop_gen2 = 1000, numgraphs = 100, plusminus_generations = 20), from 0 to 8 admixture edges.

### Species Delimitation

We conducted hierarchical heuristic species delimitation under the MSC-M model using HHSD v.0.9.8 (Kornai et al. 2024), which automates the estimation of the genealogical divergence index (GDI) in Bayesian Phylogenetics and Phylogeography (BPP). The GDI (Jackson et al. 2017) measures the probability that two alleles from the same population coalesce before either coalesces with an allele from a different population, and before the time of population divergence. Values range from 0 to 1, with values near 1 indicating strong population divergence (approaching species-level separation) and those near 0 reflecting a single, panmictic population. A meta-analysis by Jackson et al. (2017), based on data from Pinho and Hey (2010), proposed thresholds where GDI < 0.2 suggests a single species, GDI > 0.7 indicates distinct species, and intermediate values (GDI = 0.2 – 0.7) remain ambiguous.

HHSD employs a recursive, automated merge-and-split algorithm based on GDI to delineate species across extensive phylogenies. Because merging and splitting analyses require a guide tree, we used the species tree topology estimated with SVDQUARTETS as the starting point. Bidirectional migration was permitted between all possible lineage pairs, including historical migration between ancestral and sister lineages. Migration rate priors were set to a gamma distribution G (0.1, 10), corresponding to a mean of 0.01 migrants per generation.

In the merge algorithm, the software progressively groups populations into a single species, starting from the tips of the tree and moving toward the root. Merges are accepted when either of the two GDI estimates for a population pair is below 0.2, and the process continues until no further merges are possible. Conversely, delimitation begins with a single species and progressively splits lineages using the split algorithm, moving from the root toward the tips. A split is accepted if both GDI estimates exceed 0.5, and at least one exceeds 0.7. This process continues until no additional splits can be made. Following the software authors’ recommendations, we ran the merge and split pipelines without thresholds to obtain GDI estimates for all lineages.

## Results

Low-coverage sequencing generated a dataset of 162,891,718 reads. Individual read counts ranged from 294,554 to 2,685,495, averaging 1,226,849 per sample (Table S1). After processing, we identified 229,344 bi-allelic SNPs, allowing for 10% missing data. Filtering for minor allele frequency (--maf > 0.01) and thinning to 100 bp resulted in a high-quality dataset of 13,936 SNPs. PCADAPT detected and excluded 757 outlier loci, yielding a final dataset of 13,179 SNPs. For analyses including *P. axillaris* as an outgroup, the VCF file contained 13,889 SNPs, excluding 624 outliers.

### Evolutionary Relationships

SPLITSTREE allowed the recognition of the major genetic lineages among each taxonomic species based on genetic distance (Fig. 2b). Using *P. axillaris* as an outgroup to guide the interpretation of the evolutionary network, we identified four instances of strong intraspecific genetic structure resulting in one paraphyletic species (*P. guarapuavensis*) and three polyphyletic species (*P. inflata*, *P. interior*, and *P. scheideana*), each with two lineages. Regarding the *P. inflata*, Pinfl2 appeared as a sister to *P. integrifolia*, whereas *P. interior* Pteri2 was strongly associated with *P. altiplana*. The remaining species formed monophyletic groups. The network center suggested a shared ancestral lineage with potential gene flow between early-diverging groups.

The network analysis provided an initial overview of species relationships; these patterns were largely consistent across subsequent phylogenetic analyses. The monophyly of the lineages remained stable across all inferences (Fig. 1, Fig. S1), except for *P. altiplana,* for which three lineages (Palti1, Palti2, and Palti3) were recovered differently by some methods. We included this division when pruning trees in comparative analyses.

According to the generalized Robinson-Foulds distance, the trees showed varying levels of concordance (Table 1). The main inconsistencies among methods were related to support for relationships at the tree base. SVDQUARTETS and SNAPP provided the weakest support for deep branches, whereas RAxML and ASTRAL provided the strongest support. However, many relationships remained uncertain. When comparing tree topologies that included all individuals, RAxML and ASTRAL produced identical results, whereas the SNAPP tree was the most divergent. The SNAPP tree was the most discordant, followed by NJ and SVDQUARTETS, as indicated by branches that diverged along different phylogenetic paths in the DENSITYTREE (Fig. S2).

**Table 1.** Generalized Robinson-Foulds (RF) distances among phylogenetic trees inferred using different methods. The RF distance measures topological discordance between trees, with a value of 0 indicating full concordance and 1 indicating complete discordance. Distances were calculated among Neighbor-Joining (NJ), SVDQUARTETS, RAxML, ASTRAL, and SNAPP topologies, revealing substantial incongruence among methods in this recently diverged, gene flow-influenced system.

|  | SNAPP | NJ | SVDQUARTETS | RAxML | ASTRAL |
| --- | --- | --- | --- | --- | --- |
| SNAPP | 0.63 | 0.63 | 0.49 | 0.31 | 0 |
| NJ | 0.36 | 0.36 | 0.34 | 0 | 0.31 |
| SVDQUARTETS | 0.12 | 0.12 | 0 | 0.34 | 0.49 |
| RAxML | 0 | 0 | 0.12 | 0.36 | 0.36 |
| ASTRAL | 0 | 0 | 0.12 | 0.36 | 0.63 |

### Genetic Diversity and Lineage Structuration

We estimated diversity indices (Table S2) for species and independent lineages identified by phylogenetic analysis. Most private alleles were found in Pteri1, whereas Palti3 had the fewest. Except for Pgua and Pgua2, the remaining lineages exhibited positive but small inbreeding coefficients, with Preit showing the highest value (0.06).

In the PCA (Fig. 2b), the first two principal components explained approximately 20% of the total variation. They grouped individuals into four main clusters, with some overlap among lineages. Individuals of monophyletic species tended to cluster closely, whereas polyphyletic and paraphyletic species occupied different genospaces, except for *P. inflata* and *P. guarapuavensis*.

The pairwise *F*_ST_ values (Table 2) revealed varying levels of genetic structure among lineages. Pteri1 showed the highest differentiation from all other lineages, with *F*_ST_ values ranging from 0.40 (compared to Pinfl1) to 0.52 (compared to Pmant). Notably, Pteri1 and Pteri2, expected to belong to the same species, had an *F*_ST_ of 0.48. Pinfl1 exhibited low genetic differentiation from the Pgua, Pgua2, Psch, and Preit. The Pgua and Pgua2 showed a low *F*_ST_ value (0.05), as did the different lineages of *P. altiplana* (0.09–0.11).

**Table 2.**
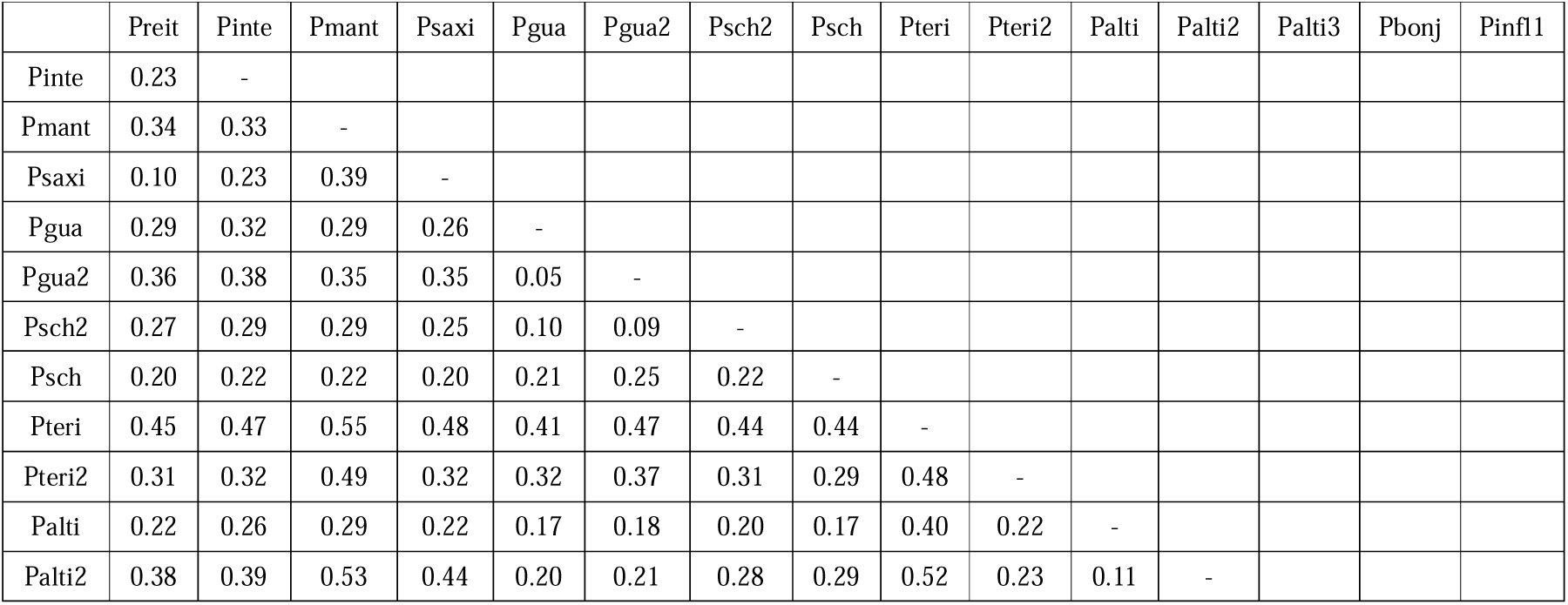
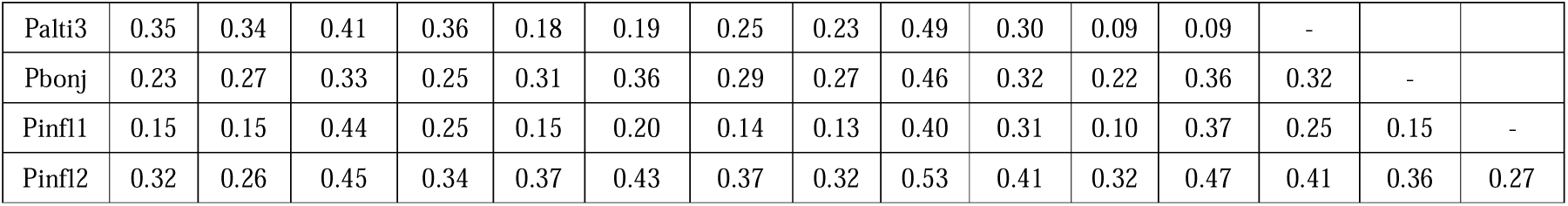
Pairwise *F*_ST_ values between lineages highlighting intraspecific genetic differentiation.

|  | Preit | Pinte | Pmant | Psaxi | Pgua | Pgua2 | Psch2 | Psch | Pteri | Pteri2 | Palti | Palti2 | Palti3 | Pbonj | Pinfl1 |
| --- | --- | --- | --- | --- | --- | --- | --- | --- | --- | --- | --- | --- | --- | --- | --- |
| Pinte | 0.23 | - |  |  |  |  |  |  |  |  |  |  |  |  |  |
| Pmant | 0.34 | 0.33 | - |  |  |  |  |  |  |  |  |  |  |  |  |
| Psaxi | 0.10 | 0.23 | 0.39 | - |  |  |  |  |  |  |  |  |  |  |  |
| Pgua | 0.29 | 0.32 | 0.29 | 0.26 | - |  |  |  |  |  |  |  |  |  |  |
| Pgua2 | 0.36 | 0.38 | 0.35 | 0.35 | 0.05 | - |  |  |  |  |  |  |  |  |  |
| Psch2 | 0.27 | 0.29 | 0.29 | 0.25 | 0.10 | 0.09 | - |  |  |  |  |  |  |  |  |
| Psch | 0.20 | 0.22 | 0.22 | 0.20 | 0.21 | 0.25 | 0.22 | - |  |  |  |  |  |  |  |
| Pteri | 0.45 | 0.47 | 0.55 | 0.48 | 0.41 | 0.47 | 0.44 | 0.44 | - |  |  |  |  |  |  |
| Pteri2 | 0.31 | 0.32 | 0.49 | 0.32 | 0.32 | 0.37 | 0.31 | 0.29 | 0.48 | - |  |  |  |  |  |
| Palti | 0.22 | 0.26 | 0.29 | 0.22 | 0.17 | 0.18 | 0.20 | 0.17 | 0.40 | 0.22 | - |  |  |  |  |
| Palti2 | 0.38 | 0.39 | 0.53 | 0.44 | 0.20 | 0.21 | 0.28 | 0.29 | 0.52 | 0.23 | 0.11 | - |  |  |  |
| Palti3 | 0.35 | 0.34 | 0.41 | 0.36 | 0.18 | 0.19 | 0.25 | 0.23 | 0.49 | 0.30 | 0.09 | 0.09 | - |  |  |
| Pbonj | 0.23 | 0.27 | 0.33 | 0.25 | 0.31 | 0.36 | 0.29 | 0.27 | 0.46 | 0.32 | 0.22 | 0.36 | 0.32 | - |  |
| Pinfl1 | 0.15 | 0.15 | 0.44 | 0.25 | 0.15 | 0.20 | 0.14 | 0.13 | 0.40 | 0.31 | 0.10 | 0.37 | 0.25 | 0.15 | - |
| Pinfl2 | 0.32 | 0.26 | 0.45 | 0.34 | 0.37 | 0.43 | 0.37 | 0.32 | 0.53 | 0.41 | 0.32 | 0.47 | 0.41 | 0.36 | 0.27 |

The remaining clustering analyses produced similar estimates of the optimal number of clusters: DAPC (Fig. 1b, Fig. S3a) identified eight groups, whereas TESS3 (Fig. 1b, Fig. S3b) and sNMF (Fig. 1, Fig. S3c) suggested nine clusters. Despite this general agreement, we observed discrepancies in the assignment of individuals to clusters across methods. As expected, most individuals were consistently grouped by putative taxonomy and geographical origin across analyses, with a few exceptions, particularly individuals from Pteri2, Pgua2, Psch2, and Psch populations, which exhibited variable assignments depending on the method used.

### Introgression/gene flow analysis

Identifying the precise path of introgression and gene flow proved to be challenging. The best introgression scheme identified by SNAQ included three events (Fig. 2b). According to this analysis, Pteri2 resulted from 15% introgression from Pbonj, with 85% of its ancestry from Palti2. Similarly, Pgua2 resulted from 21% introgression from Pgua, with 78% of its ancestry from Psch2. Finally, the clade including Pgua, Pgua2, and Psch2 resulted from a 23% introgression from the ancestral population of Pmant and Psch, with 77% of its ancestry from Pteri.

ADMIXTURE also identified three migration edges as the best number of migrations, but they were different from those inferred in SNAQ, and resulted in a very complex and reticulate pattern (Fig. 3d). In this analysis, Psch and Psch2 formed a clade that resulted from 49% introgression from Pgua2 and 51% from Pmant. The other two events included a more complex set of lineages. In the first case, a clade comprising Pbonj, Pmant, Psaxi, and Preit resulted from introgression between Pteri2 (28%) and a clade comprising both *P. guarapuavensis* lineages (72%). In the remaining event, a clade including Pteri1, Pinfl1, Pinfl2, and Pinte would have received 95% of its ancestry from the common ancestor with the clade including Pmant, Psaxi, and Preit, and 5% from the ancestral lineage of all ingroup lineages.

**Figure 3.**
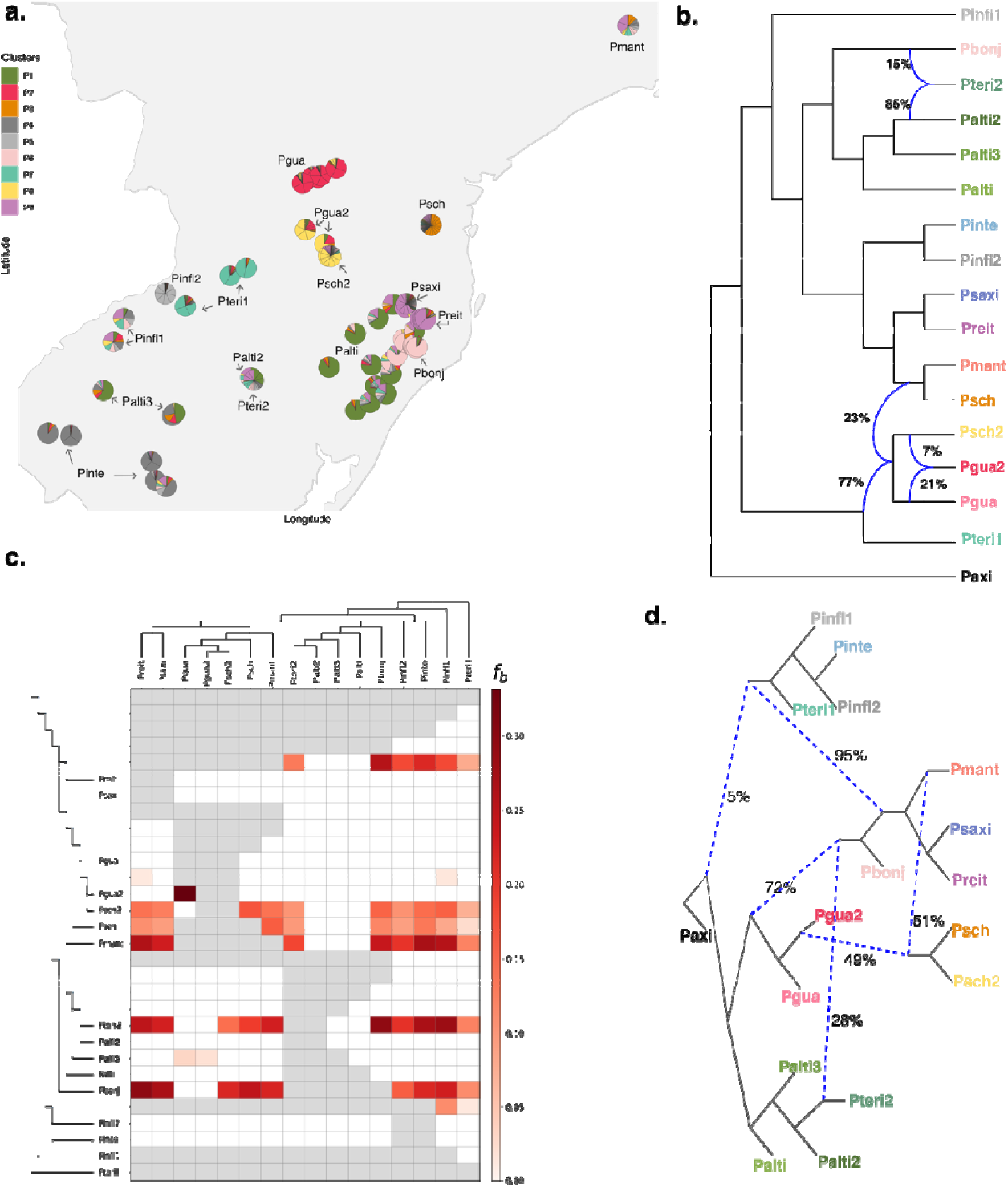
Evidence of ancestral polymorphism and widespread gene flow among highland *Petunia* lineages. (a) Geographic distribution and genetic structure based on ancestry coefficients estimated by TESS3, represented as pie charts at sampling sites. Proportions reflect individual ancestry assignments derived from genetic clustering using all SNPs. (b) Phylogenetic network inferred with SNAQ, with the best-fit model supporting three migration edges (blue branches), indicating ancestral gene flow proportions along branches. (c) *F*-branch statistics estimated with DSUITE, using the SVDQUARTETS tree topology as a reference, illustrating excess allele sharing and potential introgression events among lineages. Collectively, these analyses reveal a history of shared ancestral polymorphism and recurrent gene flow that shape the evolutionary history of these lineages. (d) Admixture graph inferred with ADMIXTOOLS2, also supporting three migration events (dashed blue lines), highlighting reticulation patterns among lineages.

The *f*-branch statistic from DSUITE revealed consistent, though slightly variable, signals of gene flow across guide trees. When the SVDQUARTETS tree (Fig. 3c) was employed as the template, several lineages (Pbonj, Pteri2, Psch1, Psch2, Pgua2, and Pmant) exhibited strong and widespread signals of introgression with nearly all other lineages, except for the *P. altiplana* lineages, which showed minimal evidence of gene exchange.

HYDE identified 4.15% of putative hybridization or introgression events (70 significant out of 1,680 analyzed trios) (Table S3). Among the parental lineages, Pinfl1 was involved in the most combinations (41 trios), followed by Pgua1 (14 trios), Pgua2 (12 trios), and Palti2 (10 trios). The lineages most frequently identified as hybrids were Pbonj (17 trios), Pteri2 (12 trios), and Pmant (10 trios). When we mapped the directionality of these events onto the phylogeny, most signals of gene flow were associated with deeper branches in the phylogeny (Fig. S4). This pattern suggests that most introgression occurred earlier in the divergence process, supporting the hypothesis of high connectivity and extensive gene exchange during the initial stages of lineage diversification. DFOIL yielded results similar to HYDE, showing a 1.68% introgression signature (2,998 introgressive out of 178,921 possible events). The most frequently involved trios were the *P. guarapuavensis* lineages with *P. reitzii*-*P. saxicola* (Table S3).

These results indicate extensive historical connectivity among lineages. Such pervasive introgression contributes to the reticulate pattern observed in the phylogenetic analyses and may obscure evolutionary relationships and species boundaries.

### Species Delimitation

Under the multispecies coalescent model (MSC-M), the HHSD split analysis did not support any lineage as distinct when using the suggested threshold of 0.7. Conversely, the merge analysis grouped all populations when the 0.2 threshold was applied. GDI estimates ranged from 0.25 to 0.86 (Fig. 1). Only three nodes, Pteri1, *P. scheideana*, and *P. mantiqueirensis*, had GDI values above 0.7, indicating strong support for their recognition as distinct species under the MSC-M (no migration) model. One additional lineage, Pinfl1, approached the threshold with a value of 0.69. All other lineages fell within the “grey zone” (0.2 < GDI < 0.7), where the boundaries among species remain ambiguous. These results highlight the challenge of delineating species from genomic data in systems characterized by recent divergence and high levels of gene flow.

## Discussion

### Incomplete Lineage Sorting (ILS) and Introgression/Gene Flow

In systems experiencing recent divergence, incomplete reproductive isolation often facilitates hybridization in zones of sympatry. At the same time, ancestral polymorphisms can persist due to incomplete lineage sorting (ILS), a process in which alleles that predate speciation are randomly allocated to descendant lineages (Maddison and Knowles 2006; Fontaine et al. 2015).

Distinguishing between these two processes remains a primary challenge in the evolutionary studies of rapid radiation, as both increase the level of shared genetic variation among lineages (Rieseberg 1997; Suh et al. 2015; Meyer et al. 2016; Edelman et al. 2019).

In this study, we show that, for *Petunia* species with short corolla tubes that inhabit high elevations, incomplete lineage sorting, gene flow, and intense introgression influenced the evolutionary relationships among species. Consequently, phylogenetic and population structure analyses cannot fully identify traditional taxonomic species as distinct isolated groups, with several lineages showing evidence of genetic admixture and shared polymorphism. Incongruences caused by ILS are particularly common when lineage divergence occurs rapidly and when ancestral populations are large (Maddison and Knowles 2006). Several studies have highlighted the relevance of shared ancestral polymorphism not only in *Petunia* (Souza et al 2022; Simon et al 2023; Pezzi et al 2024) but more generally in other groups that underwent rapid radiations (*e.g.,* Carbone et al. 2014; Wickett et al. 2014; Lamichhaney et al. 2015; Guerrero et al 2017; Naciri and Linder 2020; Olave and Meyer 2020). These shared polymorphisms can play significant roles in adaptation and speciation (Zhou et al. 2017) and have been widely reported across the Solanaceae and other plant radiations (Pease et al. 2016).

However, gene flow, either recent or ancient, is another process that can account for shared polymorphism among species. To better disentangle gene flow from ILS, we applied multiple complementary approaches to detect gene flow while accounting for its effects. All analyses consistently identified multiple instances of genomic introgression among the studied lineages (Fig 2, Fig 3). While the precise gene flow pathways depended on the species tree topology assumed, all analyses indicated that ILS was insufficient to explain the level of genealogical discordance among these *Petunia* species (Table S3, S4, Fig. S4). This conclusion is consistent with findings from other plant groups, in which hybridization and introgression have played central roles throughout the evolutionary history (Mallet 2005; Wood et al. 2009; Abbott et al. 2013; Yakimowski and Rieseberg 2014; Stull et al. 2023).

Given the extent of gene flow and genetic introgression, simple bifurcating trees do not accurately represent the evolutionary history of lineages. On the other hand, phylogenetic networks provide a more realistic framework for visualizing the complex histories by relaxing the assumption of strictly bifurcating relationships (Bastide et al. 2018; Guo et al. 2023; Steenwyk et al. 2023).

Based on our findings, we propose that a reticulate phylogeny, as shown in Fig. 2a, better represents the evolutionary relationships among the recently and rapidly diverged lineages of *Petunia* and other plant groups.

### Species Delimitation and Genome-Wide Data

Under the General Lineage Species Concept, the essential property of a species is its existence as an independently evolving lineage that maintains its evolutionary identity through time (de Queiroz 2005; 2007). Reproductive isolation, morphological or genealogical distinctiveness, and the monophyly of its subpopulations, among other characteristics, are proxies for identifying species, but, on their own, are not sufficient to delimit species boundaries (de Queiroz 2005).

Importantly, genealogical methods for delimiting species, even if based on extensive genomic data, should be seen as an additional piece of evidence for understanding what species are, but that should be interpreted in the context of morphological and ecological variation, reproductive isolation, and more. For example, the GDI metric, despite being useful, has limitations. Its results depend on effective population size and divergence time, with a broad zone of indecision (Leaché et al. 2019). Additionally, congruence among methods does not necessarily confirm accuracy if those methods share similar biases (Carstens et al. 2013; Rannala 2015).

Speciation is a gradual process. Genomic divergence occurs unevenly, influenced by selection, drift, gene flow, and mutation rates (Noor and Feder 2006; Nosil et al. 2009; Nosil and Feder 2012; Abbott et al. 2013; Zheng et al. 2017). Reproductive isolation accelerates genomic divergence by curbing gene flow between diverging populations. However, complete reproductive isolation may require a long time to establish (Coyne and Orr 2004; Marques et al. 2019) and is not always necessary for species recognition, especially in the presence of ecological divergence (Feder et al. 2012; Fujita et al. 2012; Suh et al. 2015). Moreover, when subpopulations are in allopatry, spatial isolation allows populations to quickly accumulate genetic differences driven by local adaptation, eventually leading to reproductive isolation as a byproduct (Grant 1981; Gottlieb 2004).

### Species Delimitation in **Petunia** from highlands

Considering the wide genealogical, ecological, and morphological variation exhibited by these *Petunia* lineages, lumping them into a few species based on incomplete reproductive isolation would neglect the substantial evolutionary distinctiveness of each lineage. Conversely, splitting them too early could lead to overestimating the number of species when there is insufficient evidence for an independent evolutionary trajectory. The most plausible interpretation is that we are observing recent and ongoing speciation processes, with these lineages at various stages of achieving complete evolutionary independence within the gray zone of speciation.

From a genomic point of view, different methods suggest a different optimal number of genetic clusters (Fig. 1), and only four clusters are consistent across all methods: Pgua, Pteri, Pbonj, and Preit+Psaxi. The GDI showed relatively high values (indicative of distinct species) for Pteri, Psch, and Pmant (Fig. 1). Finally, the species network (Fig. 2a) revealed that, while most lineages appeared monophyletic, notable exceptions were *P. inflata* and *P. interior*, which each formed two distinct genetic lineages. In the case of *P. inflata*, Pinfl2 appeared consistently as sister to (but distinct from) *P. integrifolia* (Pinte), whereas Pinfl1 was very distinct in the evolutionary network (Fig. 2a). Regarding *P. interior*, Pteri1 was identified as a distinct genetic lineage in virtually all analyses (Fig 1b; Fig. 2b), whereas Pteri2 tended to be closer to *P. altiplana* (in particular, Palti2) and associated with introgression events as either a descendant (Fig. 2c) or parental (Fig. 2d) lineage.

*Petunia altiplana* exhibited a wide variation (Fig. 1b; Fig. 2a, 2b; Fig. 3a), and some analyses suggested three genetic groups with a strong geographic component (Fig. 3a), which is also consistent with the genetic proximity between Palti2 and Pteri2. However, most results indicate a close relationship among all *P. altiplana* individuals, including Pteri2. *Petunia altiplana* and *P. interior* differ in the floral morphology, and individuals of each lineage displayed the morphology corresponding to their taxonomic type. We hypothesize that they share a biogeographic history.

*Petunia altiplana* western populations inhabit the lowest-elevation range of their species (Soares et al., 2024), and *P. interior* is found mainly in lowlands; the Pteri2 population lies at the edge of the species distribution, near the *P. altiplana* western area. So, we hypothesize that they were probably originally close to the ancestral population of lowland *Petunia* that colonized the highlands.

Two well-defined genealogical clusters were *P. bonjardinensis*, whose genetic distinctiveness was suggested in almost all analyses, and the group of *P. saxicola* and *P. reitzii*. This species-pair behaved as sister lineages in all analyses and formed distinct clusters in the evolutionary network (Fig. 2b). Although they are geographically very close, they occupy different ecological niches (Soares et al., 2025). Another interesting case is *P. mantiqueirensis*, which was distinct in the evolutionary network (Fig. 2b) and one of the few lineages supported as an independent entity by GDI (Fig. 1c). Despite the large geographical distance, this species was close to *P. scheideana* (especially to lineage Psch) in several analyses, and relatively close to another group composed of two distinct lineages of *P. guarapuavensis* (Pgua and Pgua2) and a lineage of *P. scheideana* (Psch2). *Petunia guarapuavensis* and *P. scheideana* were once considered synonyms due to their morphological similarities (Stehmann et al. 2009). However, genetic analyses demonstrated enough divergence to recognize *P. scheideana* as a distinct species (Soares et al, 2024; 2025). Indeed, our results showed that Psch consistently had a high GDI. However, the relationship between Pgua, Pgua2, and Psch2 is much less clear, with different analyses providing support for distinct tree topologies and introgression pathways (Fig. 1; Fig. 2). Despite most analyses suggesting a closer relationship between the geographically close Pgua2 and Psch2, how distinct these lineages are from Pgua is open to debate.

## Conclusions

Recently and rapidly diverging species challenge evolutionary inference, especially in systems characterized by pervasive ILS, gene flow, and introgression. When divergence occurs without clear reproductive or spatial barriers, isolating the direction and magnitude of gene exchange becomes increasingly difficult. Our results show that most lineages in this system are not fully independent evolutionary units but rather points along a speciation continuum. Gene flow was essential during diversification, blurring species boundaries and producing reticulate evolutionary patterns. Traditional phylogenetic trees may oversimplify relationships in such cases, while phylogenetic networks provide a more realistic representation of lineage history. Broad, integrative analyses like those applied here are essential for capturing these dynamics, which highlights the need to reassess how we should delimit species in complex, reticulate systems.

## Funding

This work was supported by Conselho Nacional de Desenvolvimento Científico e Tecnológico (CNPq) and Programa de Pós-Graduação em Genética e Biologia Molecular (PPGBM-UFRGS). CNPq and Programa Institucional de Internacionalização – Coordenação de Aperfeiçoamento de Pessoal de Nível Superior (Print-CAPES)/Universidade Federal do Rio Grande do Sul (UFRGS) supported LSS. NJRF is supported by CNPq, grant number 316900/2023–0.

## Data Availability Statement

Raw sequences are deposited in the SRA (codes available in Supplementary Table S1). Input data for all analyses are available on FigShare: 10.6084/m9.figshare.31851100.

## List of supplementary tables and figures

**Table S1**. Summary of sampled individuals and sequencing statistics. BHCB - voucher number at the BHCB herbarium (Universidade Federal de Minas Gerais, Belo Horizonte, Brazil); Long - longitude; Lat - latitude.

**Table S2**. Summary of genetic diversity indices for each lineage, including number of private alleles (Private), number of individuals (N), expected heterozygosity (Exp_Het), expected homozygosity (Exp_Hom), nucleotide diversity (π), and inbreeding coefficient (*F*_IS_).

**Table S3.** Result from the hybridization detection analysis in HYDE, only including significant results with sensible values of gamma (0 < γ < 1).

**Table S4** – Summarized result of DFOIL with the frequency of inferred migration events among lineages based on SNP data.

## Supplementary Tables available at

https://drive.google.com/drive/folders/12m2QJRahKIgqpPVMzUr_L3Eyo31AsUZl?usp=drive_link

---

**Figure S1.**
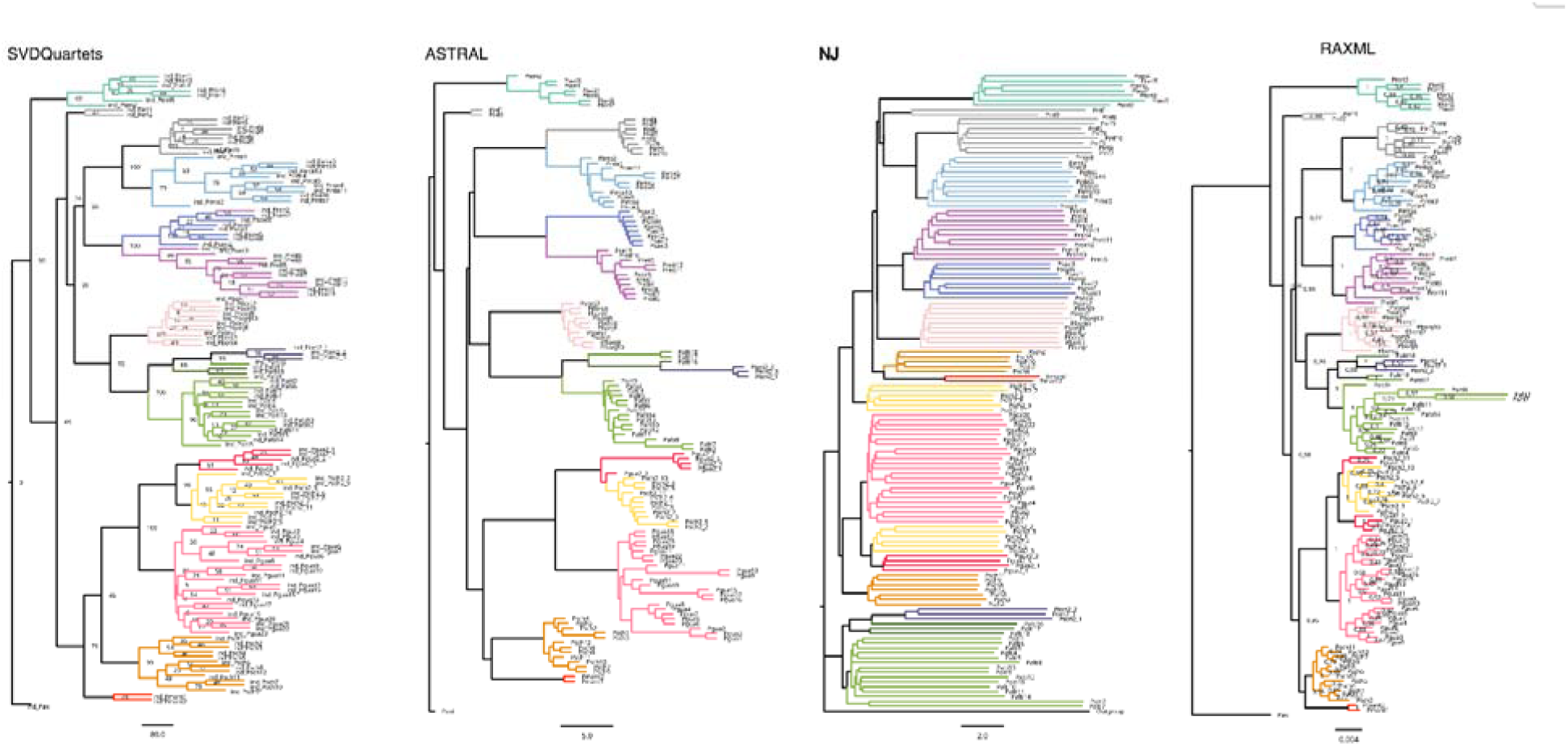
Phylogenetic trees reconstructed using different methods: SVDQUARTETS, ASTRAL, Neighbor-Joining (NJ), and RAxML. Trees are shown separately to highlight topological congruence and conflict among the methods. Lineages are colored according to the standardized scheme used throughout the study.

**Figure S2.**
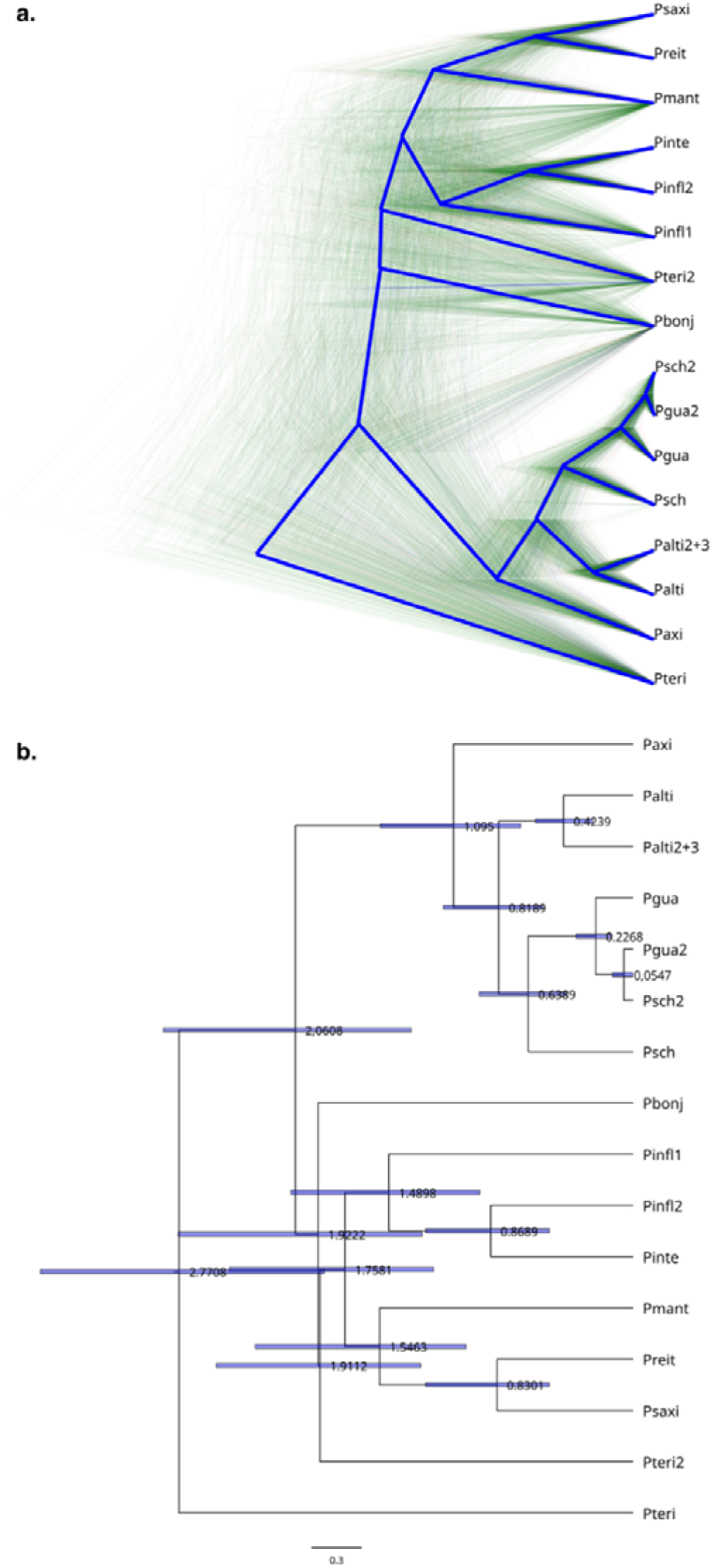
Species tree estimation based on SNAPP analysis. (a) DENSITREE cladogram displays the species tree posterior distribution, with the consensus tree highlighted in blue. (b) Consensus species tree with divergence time estimates at each node and 95% highest posterior density (HPD) intervals shown as grey bars.

**Figure S3.**
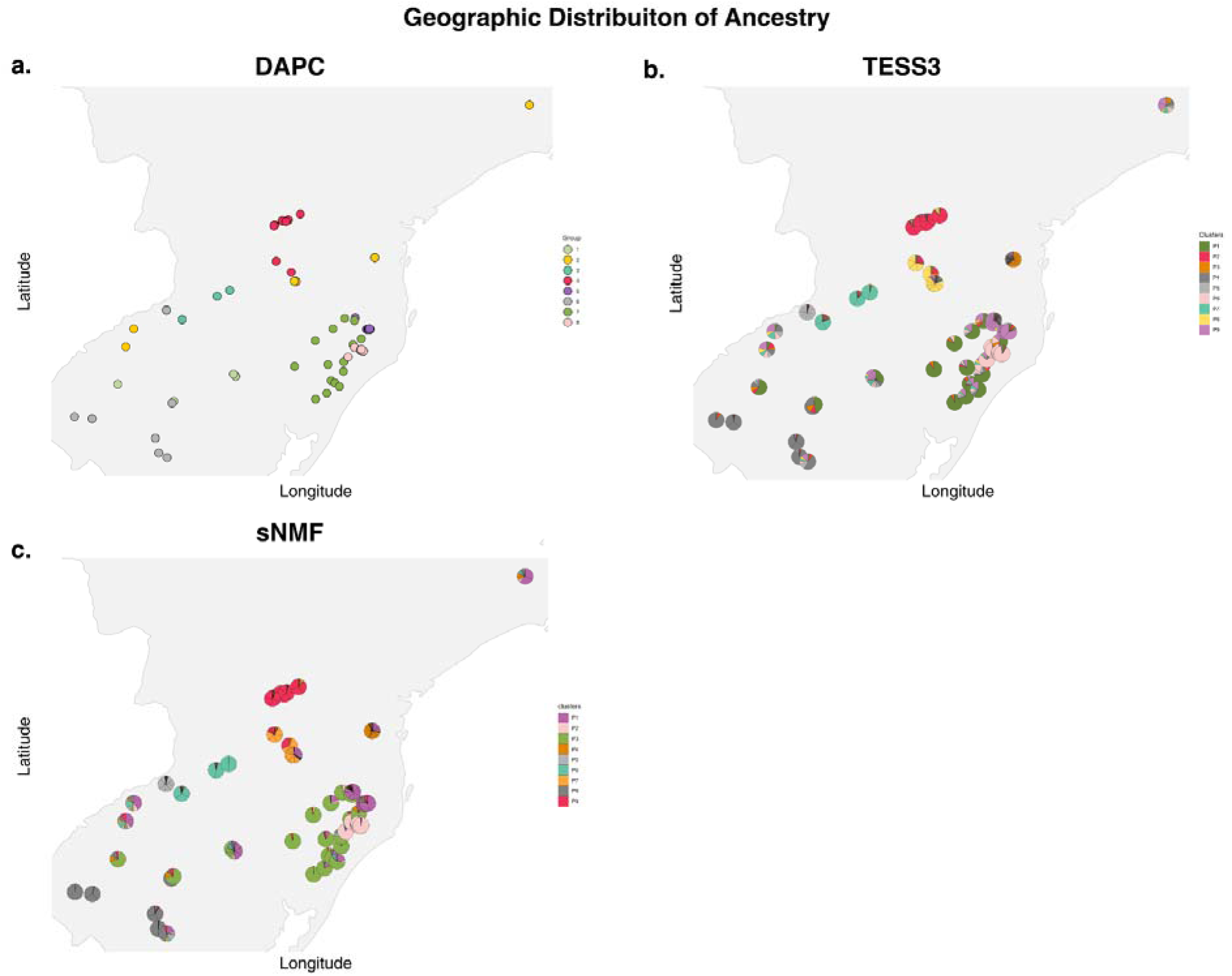
Genetic clustering results for all individuals based on SNP data. (a) Discriminant Analysis of Principal Components (DAPC). (b) TESS3 ancestry coefficients mapped for each individual. (c) sNMF ancestry coefficients. Colors are consistent with lineage assignments in other figures.

**Figure S4.**
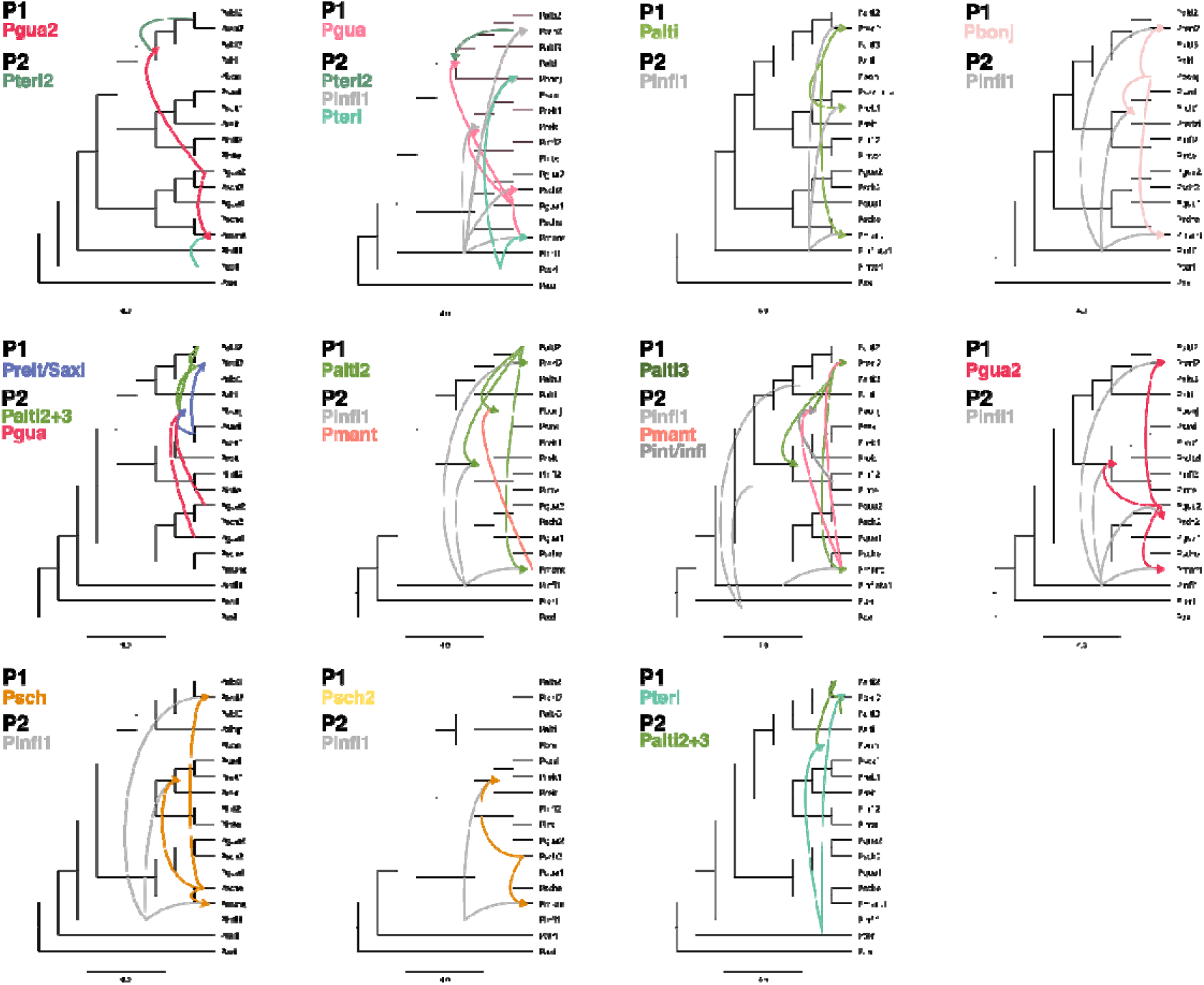
Phylogenetic trees showing inferred directions of hybridization events detected by HyDe. Arrows indicate significant hybridization signals, with colors corresponding to the inferred parental lineages. Tree topology is based on SVDQUARTETS analysis.

## Notes

https://doi.org/10.6084/m9.figshare.31851100

### Competing Interest Statement

The authors have declared no competing interest.

